# Productive lifespan and resilience rank can be predicted from on-farm first parity sensor time series but not using a common equation across farms

**DOI:** 10.1101/826099

**Authors:** I. Adriaens, N.C. Friggens, W. Ouweltjes, H. Scott, B. Aernouts, J. Statham

## Abstract

A dairy cow’s lifetime resilience and her ability to re-calve gain importance on dairy farms as they affect all aspects of the sustainability of the dairy industry. Many modern farms today have milk meters and activity sensors that accurately measure yield and activity at a high frequency for monitoring purposes. We hypothesized that these same sensors can be used for precision phenotyping of complex traits such as lifetime resilience or productive lifespan. The objective of this study was to investigate if lifetime resilience and productive lifespan of dairy cows can be predicted using sensor-derived proxies of first parity sensor data. We used a data set from 27 Belgian and British dairy farms with an automated milking system containing at least 5 years of successive measurements. All of these farms had milk meter data available, and 13 of these farms were also equipped with activity sensors. This subset was used to investigate the added value of activity meters to improve the model’s prediction accuracy. To rank cows for lifetime resilience, a score was attributed to each cow based on her number of calvings, her 305-day milk yield, her age at first calving, her calving intervals and the days in milk at the moment of culling, taking her entire lifetime into account. Next, this lifetime resilience score was used to rank the cows within their herd resulting in a lifetime resilience ranking. Based on this ranking, the cows were classified in a low (last third), moderate (middle third) or high (first third) resilience category. In total 45 biologically-sound sensor features were defined from the time-series data, including measures of variability, lactation curve shape, milk yield perturbations, activity spikes indicating estrous events and activity dynamics representing health events. These features, calculated on first lactation data, were used to predict the lifetime resilience rank and thus, the classification within the herd (low/moderate/high). Using a specific linear regression model progressively including features stepwise selected at farm level (cut-off *P*-value of 0.2), classification performances were between 35.9% and 70.0% (46.7 ± 8.0, mean ± standard deviation) for milk yield features only and between 46.7% and 84.0% (55.5 ± 12.1, mean ± standard deviation) for lactation and activity features together. This is respectively 13.7 and 22.2% higher than what random classification would give. Moreover, using these individual farm models, only 3.5% and 2.3% of the cows were classified high while being low and vice versa, while respectively 91.8% and 94.1% of the wrongly classified animals were predicted in an adjacent category. A common equation across farms to predict this rank could not be found, which demonstrates the variability in culling and management strategies across farms and within farms over time. The lack of a common model structure across farms suggests the need to consider local (and evidence based) culling management rules when developing decision support tools for dairy farms. With this study we showed the potential of precision phenotyping of complex traits based on biologically meaningful features derived from readily available sensor data. We conclude that first lactation milk and activity sensor data have the potential to predict cows’ lifetime resilience rankings within farms but that consistency over farms is currently lacking.

## INTRODUCTION

Increasing the longevity of dairy cows is key for the dairy sector’s sustainability. Cows with a long productive lifespan typically have a good reproductive performance, few health problems and produce milk in an efficient and consistent way. A dairy cow typically only starts to make profit for the farmer during her second lactation and she reaches her full production potential as late as in her third lactation (Cabrera, 2018). Early culling and short longevity thus clearly has a negative impact on the economic efficiency of the herd. More importantly, longevity and optimizing the ratio between the productive and non-productive life is also crucial for the fulfillment of societal demands and to reduce environmental impacts of the sector (van Knegsel et al., 2014).

One step towards the optimization of the farm management with respect to longevity would be the identification of animals that have a high probability of completing several lactations, or more specifically, that are ‘resilient’. Resilient animals can be considered as animals that avoid early culling by coping well with the farm’s management conditions. These animals reproduce easily, produce consistently and react well on imposed challenges and (physiological) stress (Ahlman et al., 2011). Correct and timely identification of these resilient animals would not only allow for optimization of culling and (advanced) breeding decisions, but also for the long-term selection of animals that thrive in their specific farm environments (e.g., infection pressure, feed quality, milking conditions, …).

Optimized efficiency on farm would require that the lifetime resilience of an animal can be predicted as soon as possible. Genetic indicators for lifetime resilience are not yet available as phenotypic information on this complex trait is lacking. Nevertheless, recent technological developments led to an increased implementation of sensor systems and automation to improve the herd management and reduce labor requirements (Steeneveld and Hogeveen, 2015). Besides for the detection of health problem and fertility events, many of these sensor systems also have the potential to provide targeted information about other, more complex traits (Friggens and Thorup, 2015). In this study, we hypothesized that common sensor data such as milk production and activity time series can be used to predict a complex trait such as lifetime resilience. Simultaneously, additional benefits of these technologies will be generated from the calculation of precision phenotypes and their use for the characterization of overall and relative performance of the animals within the farm context and compared to herd mates (Royal et al., 2000; Tenghe et al., 2015; Sorg et al., 2017). Accordingly, when sensor data can be used to this purpose, selection of animals on these more complex traits becomes possible, which, when combined with the genetic merit of each animal, can boost future breeding efforts at farm and population level (Köning and May, 2019).

In order to use sensor data for the prediction of lifetime resilience such that it can be used for both decision support and precision phenotyping, we propose to derive biologically meaningful proxies for the cow’s physiological status from the high-frequency milk yield and activity dynamics available from commercially available sensor systems. It was shown previously that each change in feed intake or energy allocation (for e.g. an immune response) may result in yield perturbations (Ben Abdelkrim et al., 2019) and thus, the milk yield dynamics mirror the animal’s physiological status. Similarly, activity dynamics reflect potential estrus and the more general behavioral responses of the cows to physiological and environmental stress (Rutten et al., 2013). Because of the link between health and fertility performance and longevity and culling, the proposed concept is that through characterization of these dynamics, it will be possible to predict their lifetime resilience.

This study aimed at developing meaningful milk yield and activity features from first lactation sensor time series and combining these features into a model capable of predicting the lifetime resilience of the cows on each farm. This model could be used to help farmers to identify animals that cope well with their specific farm contexts, for example to aid breeding (e.g., dam selection for sexed semen, embryo transfer or the use of a beef sire) or culling decisions as early as after the first lactation. As such, there is still time to take decisions that directly contribute to the farm’s efficiency by selecting animals which perform well on that particular farm.

## MATERIALS AND METHODS

### Data Collection and Selection

#### Available Data

Software back-ups of the farm management system were collected on respectively 34 and 42 Belgian and British farms with an automated milking system (**AMS**). From this database, 27 farms were selected based on (1) the accessibility and reliability of at least 5 years of contiguous data, and (2) the availability of daily milk yield at individual cow level. The time period covered by these data varied between 2005 and 2019. All these 27 farms had either an AMS of Lely (Lely Industries N.V., Maasluis, the Netherlands; No. = 16) or of DeLaval (DeLaval International, Tumba, Sweden; No. = 11). On average 2.4 lactations were recorded per cow’s life. All farms had intensive production systems with cows kept indoors and fed with both forage and concentrates. Other management practices differed among herds but were not further documented in the software back-up files of the farm management system.

All data tables were extracted from the restored back-up files of the AMS software system using SQL Server Management Studio (Microsoft, Redmond, WA, United States). The further data mining, pre-processing and merging of these data tables and the rest of the analyses described below was done in Matlab R2017a (The MathWorks Inc., Natick, MA, United States). Both the full data set of 27 farms all having daily milk records (data set 1, **DS1**) and a subset of 13 farms also having daily activity data available (data set 2, **DS2**, all milked by a Lely AMS) were used for this study. An overview of the characteristics of both data sets is given in Table 1.

**Table 1.** Overview of the available data sets. Data set 2 (**DS2**) is a subset of data set 1 (**DS1**) for which also daily activity data were available besides milk meter data.

| Descriptor | Data set 1 (DS1)<br>Average ± SD [range] | Data set 2 (DS2)<br>Average ± SD [range] |
| --- | --- | --- |
| No. of days data per farm | 3001 ± 828 [1830 ; 4977] | 3020 ± 815 [1837 ; 4101] |
| No. of cows in total | 3754 | 2075 |
| No. of cows per farm | 139 ± 82 [24 ; 308] | 160 ± 94 [57 ; 308] |
| No. of lactations in total | 9395 | 5286 |
| No. of lactations per farm | 348 ± 229 [44 ; 799] | 407 ± 264 [113 ; 799] |
| Age at first calving per farm (years) | 2.2 ± 0.1 [2.0 ; 2.4] | 2.2 ± 0.1 [2.0 ; 2.3] |
| Average 305-day milk yield per farm (kg) | 9557 ± 1007 [7931 ; 11739] | 9691 ± 1061 [8244 ; 11739] |
| Average calving interval per farm (days) | 407 ± 16 [377 ; 433] | 403 ± 15 [377 ; 432] |
| Average dry period length per farm (days) | 56 ± 13 [35 ; 91] | 59 ± 16 [35 ; 91] |

#### Cow Selection

After extraction of data tables, individual cows on each farm were selected based on the availability of sensor data for their entire lifetime production. Because exact culling dates were not always available, we elected to apply a criterion to discriminate between cows that likely had been dried off towards the end of the time span covered by the dataset available for that farm (not to be included in the analysis) or removed from the herd (to be included in the analysis). For each farm, the 95% confidence interval on the average dry period length was calculated. If the last milk record was before the end of the data set minus the upper 95% confidence interval boundary, that cow had 97.5% chance that she was removed from the herd and she was included. Accordingly, only cows that met this criterion and for which the date of her first calving was within the timespan of the available data for that farm were selected. An overview of the characteristics of this selection is provided in Table 1. Data set 1 consisted of 3754 unique cows and 9395 unique lactations, while DS2 included 2075 cows with 5286 lactations. Per farm, respectively 24 to 308 cows (139 ± 82, mean ± SD) with in total 44 to 799 lactations (348 ± 229) and 57 to 308 cows (160 ± 84) with 113 to 799 lactations (407 ± 264) were selected for DS1 and DS2.

#### Sensor data

The milk yield sensor data were recorded by the AMS using ICAR-approved milk meters as integrated in the Lely and DeLaval robots. The available activity sensor data were recorded by Lely (Lely Industries N.V., Maasluis, the Netherlands) neck-mounted activity sensors and consisted of raw 2-hourly measures of acceleration in x, y and z planes that were aggregated in single daily activity records, but no further details of the individual sensor systems were available. Although in this study the raw data were all similar with 2-hourly values varying between 0 and 300, the presented methodology is independent of the actual raw values and can be applied on all measures of activity for which daily aggregates are available as long as they represent the behavioral changes of the cows linked with estrous and health events.

### Calculation of Lifetime Resilience Ranking

For this study, the lifetime resilience of a cow was considered primarily as the cumulative result of her ability to recalve (and thus, to extend her productive lifespan) supplemented with secondary corrections for age at first calving, calving intervals, 305-day milk yield, health events and number of inseminations (Friggens and De Haas, 2019). With a high weight given to each newly started lactation (i.e. parity number), the secondary corrections mainly allow discrimination between all cows reaching a certain parity. This definition was agreed upon within the EU Horizon 2020 GenTORE consortium consisting of researchers, animal experts, veterinarians, technology suppliers and geneticists and is further detailed in Friggens and De Haas, (2019). Because the number of inseminations and health events were not consistently available for all herds over the entire time period, the final equation for calculating lifetime resilience scores (**RS**) excluded these variables and was (Eq. 1):

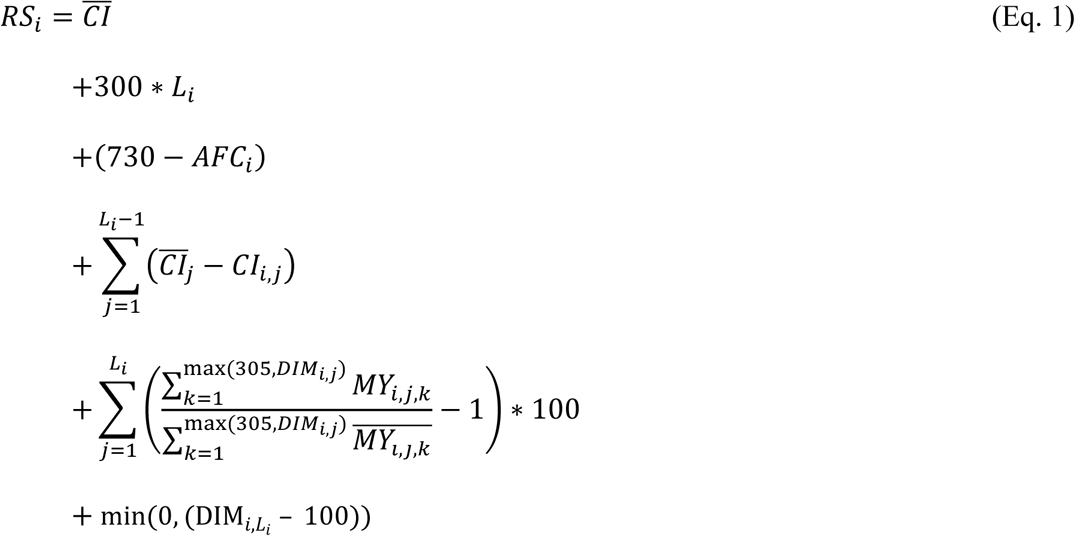

With:

*RS*_*i*_ Lifetime resilience score for cow *i*

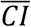 Average calving interval of the herd

*L*_*i*_ Lactation number in which cow *i* exited the herd (last lactation number of a cow)

*AFC*_*i*_ Age at first calving of cow *i* in days

*CI*_*i,j*_ Calving interval of cow *i* between the start of lactation *j* and (*j* + 1)

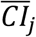 Average calving interval between the start of lactation *j* and (*j+1*) of all cows in the herd

*MY*_*i,j,k*_ Milk production (in kg) of cow *i* at day *k* of lactation *j*

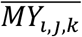 Average milk production (in kg) at day *k* of all cows in the herd in lactation *j*

*DIM*_*i,j*_ Days in milk (**DIM**) of cow *i* at the end of her lactation *j*

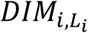 Days in milk of cow *i* at the end of her last lactation *Li*

This way, each RS consists of (1) a baseline equal to the average calving interval of that herd to avoid negative lifetime resilience scores (this does not contribute to the ranking); (2) a bonus of 300 points given for each recalving (newly started lactation); (3) a penalty or bonus score given to cows older respectively younger than 24 months at their first calving equal to 1 point per day longer or shorter than 730 days (i.e. 24 months); (4) a penalty or bonus score equal to the number of days the calving interval is respectively shorter or longer than the average calving interval of the same parity in the herd; (5) a penalty or bonus score equal to the percentage the 305-day milk production is respectively lower or higher compared to the average 305-day production of the corresponding parity for all lactations in the herd, reflecting production performance in the most relevant part of the lactation; and (6) a penalty score equal to 100-*DIM*_*exit*_ for cows exiting the herd before day 100 in lactation assuming that these cows are involuntarily removed from the herd.

Based on this score, which assumes that these factors reflect the accumulated effects of the cows’ resilience over their lifetime, cows were ranked within farms. For example, if the average CI of a herd is 400 days, the average CI between the start of first and second parity is 380 days and the average 305-day milk production in first lactation is 8000 kg, then a cow that calved twice (the first time aged 775 days), had a calving interval of 420 days between the start of the first and second lactation, produced 5% more milk than average in the first 305 days of the first lactation (8400 kg) and 20% less than her herd peers in the first 90 days of the second lactation, and that was culled day 90 in the second lactation will receive a lifetime resilience score of RS = 400 + 300 * 2 + (730 – 775) + (380 – 420) + 5 – 20 + (90 – 100) = 890 points. Because the weights in the lifetime RS were arbitrarily chosen and based on expert knowledge, and the main interest is to distinct the lowest from the highest resilient animals, the RS was converted into an on-farm resilience rank (**RR**) to rank the cows for their lifetime resilience performance. This way, the exact weights or points assigned to each variable in Eq. 1 are of less importance. In the final lifetime RR, high ranked cows (‘highly resilient animals’) represent animals recalving many times, having the (theoretically) optimal age at first calving, having short calving intervals (and thus good reproductive performance), and producing proportionally more milk compared to their herd mates. The number of lactations affects this ranking the most because of the high weight for each new lactation started. Before entering the lifetime RR in the models, it was scaled for each farm using *RR*_*scaled*_ = (*RR*-*RR*_*min*_)/(*RR*_*max*_ – *RR*_*min*_) with *RR*_*min*_ equals 1 and *RR*_*max*_ equals the maximum rank (equivalent to the number of cows included in the ranking for that farm). The resulting *RR*_*scaled*_ varied between 0 (i.e. the highest ranked cow) and 1 (i.e. the lowest ranked cow), and thus, no scale effects caused by the varying number of animals included per farm would influence the prediction models. In the rest of this manuscript, the RR refers to the scaled RR.

### Sensor Features (SF)

#### Milk Yield SF

The time-series data of two sensors were included in this study: (1) milk meter sensors from which daily milk yields were calculated and (2) activity sensors from which the two-hourly raw data were aggregated into daily activity records. Sensor features were calculated for each of the cows for which the first lactation was longer than 200 days, because 200 days is enough to grasp a good image of the time-series dynamics as the second part of the lactation curve after the peak can be estimated by a linear function (Wood, 1967). Only the data of the first 305 days of the first lactation were included for the calculations.

As explained more in detail in Appendix A, in total 30 milk yield SF were calculated based on the daily milk yield dynamics. Moreover, a methodology was developed to calculate SF from the dynamics of the lactation curves using both the theoretical shape and the deviations from this theoretical shape as proxies for the cow’s physiological status. Accordingly, SF were defined in the following categories: (1) lactation shape characteristics including peak yield, consistency, days in milk of peak, etc.; (2) goodness-of-fit and variability measures including the characteristics of lactation model residuals; and (3) perturbation features characterizing the disturbances in the lactation dynamics and including the development and recovery rates, the number of perturbations, etc. To determine the theoretical shape of the lactation curve (i.e., potential production when no perturbations are present), a simple lactation model was iteratively fitted such that perturbations were excluded and thus, did not influence the lactation model’s coefficients (Adriaens et al., 2018). The chosen model was the nonlinear Wood model: ***TMY*** *= A*e*^*-B*DIM*^*\*DIM*^*C*^ with *A, B* and *C* being the model’s coefficients, *TMY* the total daily milk yield in kg and *DIM* the days in milk expressed in days (Wood, 1967). The Wood model (gamma function) describes the lactation curve with an increasing phase, a peak and an almost linear decreasing phase, describing the overall lactation dynamics with only three coefficients. Its simplicity reduces the computational power needed when repeatedly fitting the nonlinear model, and therefor this equation was preferred over more complex lactation models. In each iteration, the residuals were calculated by subtracting the fitted Wood model from the milk yield data. Next, all the residuals smaller than 85% of the theoretical curve (i.e. Wood’s model) were removed and the model was refitted in a next iteration. This procedure was repeated for each lactation curve individually until the difference of the average root mean squared error (**RMSE**) between two iterations was smaller than 0.10 kg for that curve, or for at most 20 iterations. The final model coefficients represent the lactation shape when no perturbations would have been present, and the model’s residuals reflect the perturbations and ‘unexpected’ milk yield dynamics. An example of the daily milk yield data, the iterated Wood model and the corresponding residuals is shown in Figure 1.

Next, lactation SF were calculated from the final coefficients of Wood’s model (*A, B* and *C*), the residuals of all daily milk yield records and the periods identified as perturbations. For the latter, major events (i.e. periods of at least 10 days of successively negative residuals with at least one day of milk production lower than 80% of the theoretical production) were discriminated from minor events (i.e. periods of at least 5 days of successively negative residuals with at least one day of milk production between 90% and 80% of the expected production). It was assumed that large perturbations (major events) may represent severe health problems which might influence culling and rebreeding decisions, while smaller (minor) perturbations are more probably linked to chronical or subclinical infections, with a different effect on culling or longevity. A detailed description of the milk yield SF and how they were calculated can be found in Appendix A.

**Figure 1.**
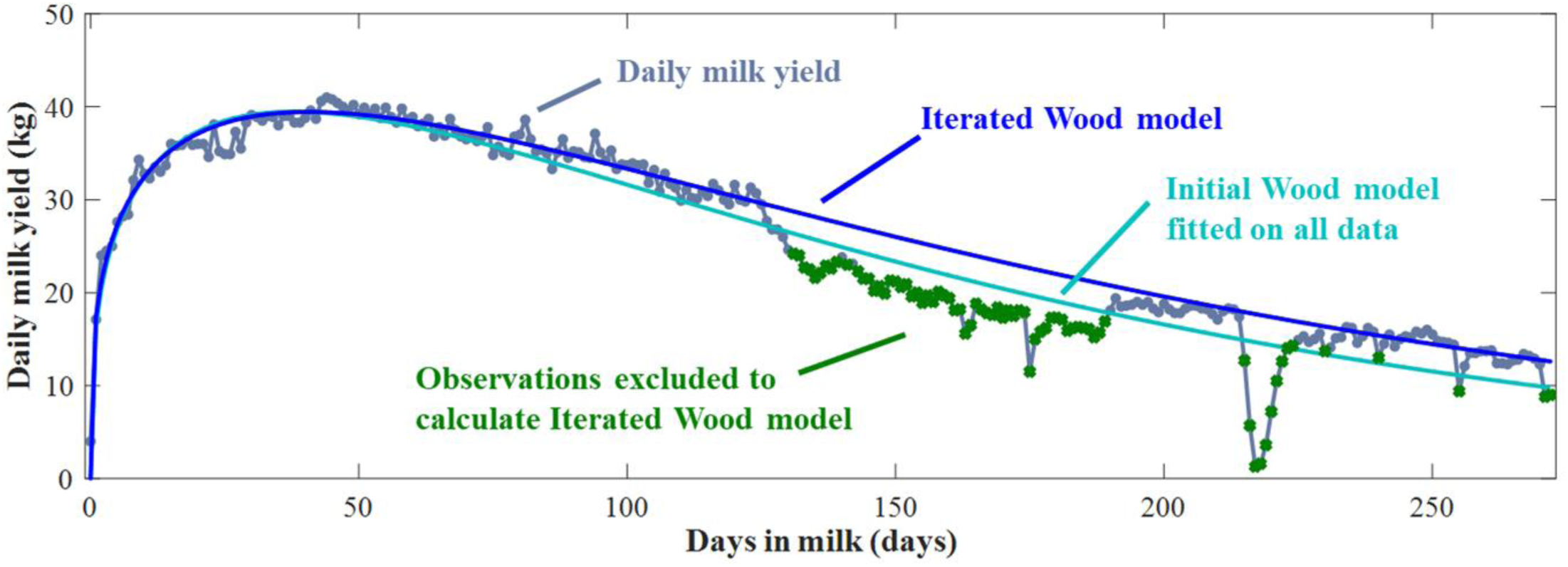
Example of a lactation curve with daily milk yields, the corresponding initial Wood model fitted on all daily milk yield data and the final Wood model fitted iteratively by excluding daily milk yields lower than 85% of the estimated curve. The residuals of the latter curve are used to characterize perturbations (e.g., representing health events).

#### Activity Sensor Features

In addition to the lactation SF, for DS2 also activity SF were calculated from the daily aggregated raw activity measures. Fifteen different SF were defined in the following categories: (1) features related to the absolute (within-herd) levels, i.e. variability and autocorrelation; (2) fertility-related characteristics based on short spikes representing estrous behavior; and (3) overall activity-related characteristics based on changes in average activity during longer periods of time which possibly relates to e.g. health events. To identify the short spikes of the second category, a median smoother using a window of 4 days was used and subtracted from the raw daily activity data to obtain residual activity levels. A short spike was identified as an increase above 40% of the maximal residual. For the identification of the longer-term patterns in the data, a 20 day-window median smoother was applied and subtracted from the daily activity data to obtain the residuals. A threshold of 20% of these minimal and maximal activity residuals was set to identify changes in activity of several days compared to the previous period. The details for the activity SF calculations are given in Appendix B. Although all the raw values in DS2 had similar variability and magnitude (i.e., with 2-hourly measures between 0 and 300 and similar longitudinal patterns), the developed procedure can be applied on all sorts of activity data, also those originating from other types of sensors independently of the exact activity levels, as long as daily measurements are available.

#### Standardization of the SF

The mean and SD differed across SF. For example, days in milk of peak varied between 10 and 150 days, while the lactation persistency comprised values between 0.001 and 1. When using the SF as variables in a prediction model, this can cause an imbalance in the fitting and selection procedure and it limits the interpretation of the model regression coefficients. Therefore, standardization of the SF was needed. Before entering the SF in the models, each SF was standardized within herd using mean centering (i.e., subtracting the within-herd mean of that SF) and by dividing them by the within-herd SD. Accordingly, the standardized SF within a herd have a mean of zero and a SD of one, which thus corrects for differences in their order of magnitude and solves interpretability issues. This standardization step ensures that a higher absolute value of a model coefficient indicates a larger effect on the model outcome (i.e., the lifetime RR). Standardized SF (both milk yield and activity) with values smaller than −3 or higher than 3 (mean ± 3*SD) were considered as outliers and replaced by zeros (i.e., the average value) to avoid missing and unbalanced data. The objective was to develop a tool to evaluate and forecast the (phenotypic) performance of an animal in the herd early in her productive life to still have time to take breeding decisions that would directly contribute to the farms’ performance, and so that ‘high risk’ animals could be monitored closer. Therefore, in this study only the first parity SF were taken into account as proxies for performance, health and fertility to predict their lifetime resilience and recalving ability on farm.

### Exploratory Analysis

In this study, a model was sought to predict the lifetime RR of all the animals on a specific farm. Ideally, a common model structure that is valid for all farms would be obtained, as this would allow the calculation of a limited and universal number of SF indicative for the animals’ lifetime resilience. As a first step to evaluate the consistency between the SF and the lifetime RR across farms, the Pearson linear correlation coefficient between each SF and the lifetime RR at individual farm level was calculated. High positive and negative correlations would indicate a strong effect of that SF on the lifetime RR, and thus a potential candidate for inclusion in further prediction models.

In a second step, mutual correlations between the SF were explored for all farms together. This initial data exploration using data of all farms together pointed out some significant (but small) linear correlations between the SF. However, at individual farm level, these correlations were often inconsistent and the sign of the correlations differed between farms. To investigate whether an underlying latent structure existed in the SF and avoid future multi-collinearity in the prediction models, a principal component analysis was carried out on the SF of both DS1 and DS2. These principal component analyses showed that respectively 8 and 24 principal components with eigenvalues higher than 1 (Kaiser criterion) explained only 71% and 74% of the variance, suggesting that a latent structure for data reduction over all farms did not exist.

### Individual Farm Model Development

Several multivariate modelling techniques including partial least squares and general linear mixed models were tested, but all had poor prediction performance (with classification results equal or worse than what random classification would have given, i.e., less than a third correctly classified) or showed significant overfitting of the data. Ultimately, a separate multivariate linear regression model relating the SF to the lifetime RR within farm was constructed as follows (Eq. 2):

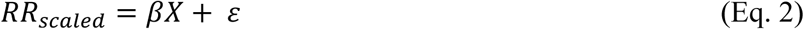

With *RR*_*scaled*_ being the scaled lifetime RR between 0 and 1 as defined above. The *β* vector contains the regression coefficients for the standardized SF in the design matrix *X*, and *ε* are the residual errors. A backward stepwise regression procedure was used to identify redundant SF in *X* applying a *P-*value of 0.2 as the inclusion threshold. This means that if there was a probability of more than 20% that removing the SF had a significant deteriorating effect on the prediction model, it was included. The chosen threshold might seem uncommonly high but given the high variability in the SF both between and within farms, we deemed relevant to include any feature having a tendency towards significance.

Ten-fold cross-validation (**CV**) was performed to evaluate the prediction performance of the obtained models and identify overfitting. To this end, ten times all the cows of each farm were assigned to either the calibration (90% of the animals) or validation (10% of the animals) set using random sampling from a uniform distribution, but applying the additional criterion that both the calibration and the validation set contained at least one animal ranked in the highest third, one in the middle third and one in the lowest third of the ranked cows. In each CV-cycle, the cows in the calibration set were used to estimate the regression coefficients *β* and the obtained model was used to predict the *RR*_*scaled*_ of the cows in the validation set. The average prediction results over all ten CV-cycles were considered to represent the final model performance.

### Model Evaluation

The initial model fit at farm level was evaluated using the RMSE (i.e., the RMSE_TR_) and the *R*^*2*^_*adj*_, calculated as shown in equation 3 (Eq. 3):

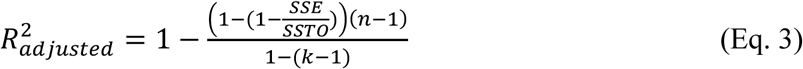

With *SSE* = residual sum of squares of the regression, SSTO the total sum of squares (i.e. the mean value of the outcome *RR*_*scaled*_), *n* the number of data points of each farm and *k* the number of SF retained in the final model for that farm.

To evaluate classification performance and discriminate between high and low resilient cows (which is of practical relevance), the cows of each farm were divided into three different categories based on their ranking: high (**H**; top third), moderate (**M**; middle third) and low (**L**, bottom third) resilient animals. When the predicted *RR*_*scaled*_ did not cover the full range of 0 to 1, and to be able to calculate these high, medium and low ranked categories for each farm, a farm-individual correction factor was applied on the predicted *RR*_*scaled*_ scores as follows (Eq. 4):

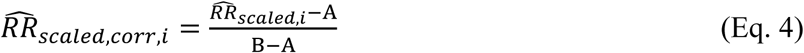

With 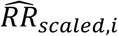 being the predicted *RR*_*scaled*_ of the *i*^*th*^ cow and *A* and *B* are farm-specific coefficients representing respectively the minimum and maximum of all the predicted *RR*_*scaled*_ for that farm in the calibration set of each CV-cycle. The *RR*_*scaled*_ of the cows in the validation set of each CV-cycle were predicted using each individual farm model (Eq. 2) and their category (H, M, L) was determined after applying the correction using the farm-specific coefficients (Eq. 4). Both the root mean squared error of cross-validation (**RMSECV**, Eq. 5) and the classification accuracy were evaluated in this CV to assess the models’ prediction performance.

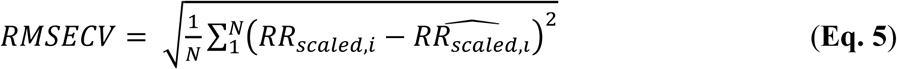

To evaluate whether a common model structure across farms could be identified or whether specific features are highly correlated to lifetime resilience in all farms, we evaluated the overlap in retained features for each farm, both in terms of their inclusion or exclusion in each farm-specific model and the sign of their regression coefficients.

Prediction performance improvement of the models including and excluding activity features was assessed using a one-sided paired *t*-test on the percentage correctly classified using the null hypothesis “*activity features do not improve (i.e. increase) the percentage correctly classified animals*” and on the proportion oppositely classified using testing the null hypothesis “*activity features do not improve (i.e. decrease) the percentage oppositely classified animals”*.

## RESULTS AND DISCUSSION

### Lifetime Resilience

This study investigates the possibility to predict ‘lifetime resilience’ from SF of the first lactation. Currently, there is no consensus on what ‘resilience’ exactly is, and definitions found in literature include the ‘ability to maintain performance regardless of pathogen burden’ and ‘the adaptation ability to a broad range of environmental conditions’ (Mulder and Rashidi, 2017; Köning and May, 2018). Our approach differs from these definitions and considers lifetime resilience as the cumulative effect of good health and fertility and a high adaptability to challenges, resulting in long productive life spans.

Figure 2 shows the lifetime resilience score plotted against the lactation in which the cows were culled for one farm as an example. Each circle represents one animal on the farm, and a higher resilience score also means a higher lifetime resilience ranking. The animals on this example farm exited the farm in between their first and seventh lactation. This figure confirms that in general for our definition of resilience, the total number of lactations each cow has started has the biggest impact on the final lifetime RR. The cows ranked lower than their herd mates with higher last lactation numbers are seen as spikes. These are mainly cows that are removed from the herd immediately after calving, and thus for which the penalty given for exiting before day 100 in lactation has a large effect (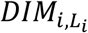 in Eq. 1).

**Figure 2.**
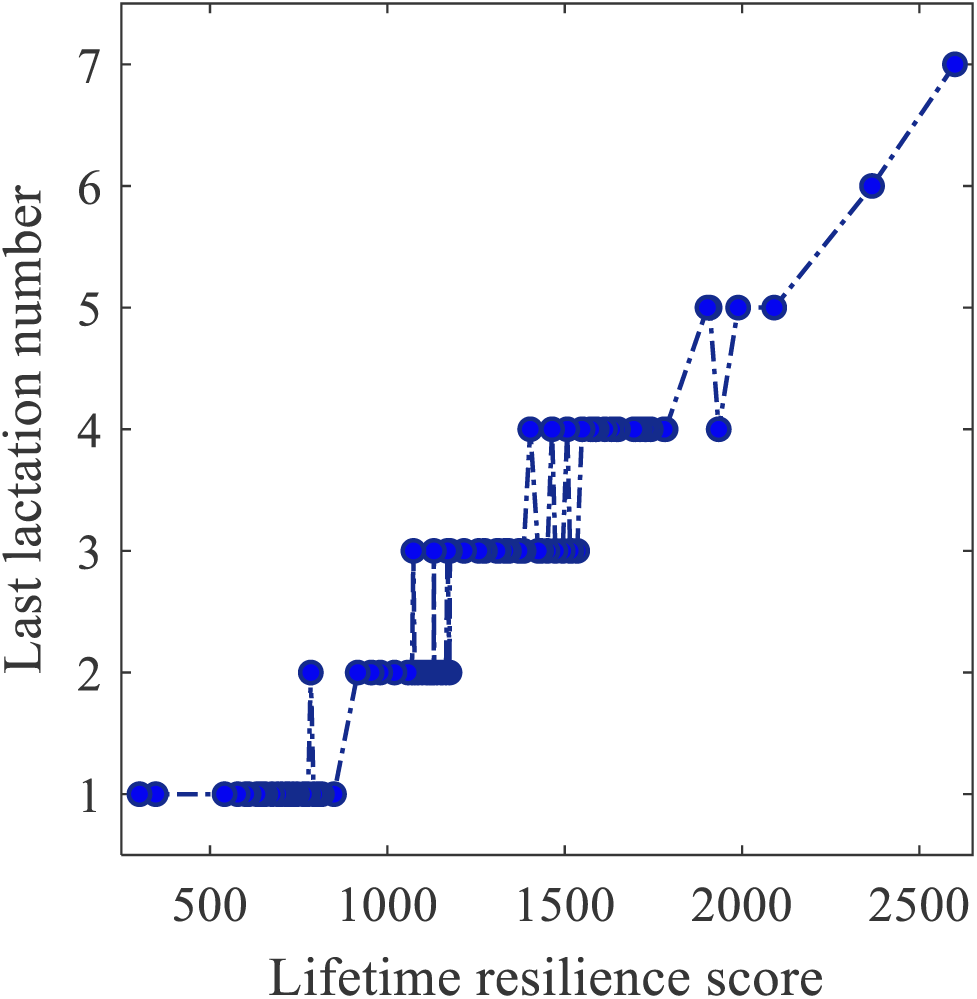
Example of the relation between the lifetime resilience scores of all the animals on an example farm (No. of cows = 110) against the parity number in which each animal exits the herd. Higher resilience scores generally correspond to higher last lactation numbers.

The inclusion of the 305-day yield originates from the idea that to differentiate two animals exiting the herd after the same number of lactations, the one with the highest production in the first 305 days (and thus, probably not encountering severe health events) would probably be the more resilient one. Ideally penalties or bonus points for health events and the number of inseminations would also be included, but the health and insemination records were not sufficiently complete over the whole time period for all farms. Consistent and correct registration, collection, mining and storage of data remains a challenge in the development of on-farm applications (Hudson et al., 2018).

### Sensor Feature Definition and Overview

The present study shows an example of how real-farm high-frequency sensor data can be used beyond monitoring and detection applications (Boichard and Brochard, 2012). We used the longitudinal, high-granularity milk yield and activity data of respectively 27 and 13 commercial dairy operations with an AMS to calculate biologically meaningful features of the cows. This is a unique data set, not only because of its commercial nature (as opposed to research farm data), but also because these high-frequency time series allowed for the inclusion of the *dynamics* of milk yield and daily activity. The availability of at least 5 years of successive measurements per farm uniquely permitted us to study the accumulated effect of health and fertility traits of many animals over their entire lifetime.

Today, cows exit the herd for many different reasons, for which the most common are poor reproduction performance, udder health problems, metabolic disorders in early lactation and claw health and locomotion disorders (Ahlman et al., 2011; Santos et al., 2016). The SF were calculated starting from expert knowledge and biological hypotheses on the supposed effect of these culling reasons on the sensor time series, and included features characterizing the perturbations and dynamics of the lactation and activity curves. For example Elgersma et al., (2018) showed that fluctuations in milk yield can reflect the cow’s health status. Table 2 gives a summary of the most important SF over all farms for DS1 and DS2. As can be seen from the minimum and maximum value of each SF, some of these SF have extreme values which upon further investigation appear to result from erroneous calculations rather than from real deviating curves. These outliers were set to zero for the analysis.

**Table 2.**
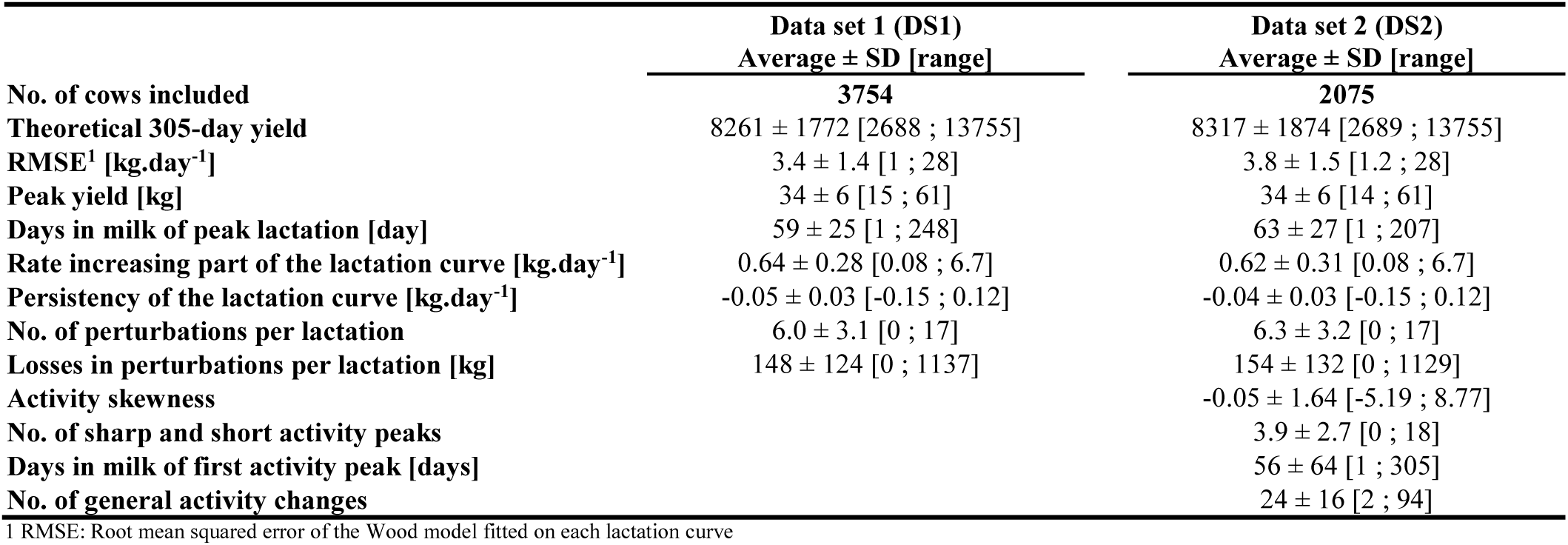
Overview of a selection of lactation and activity sensor features calculated on first parity data for data set 1 (**DS1**) and data set 2 (**DS2**). A detailed overview for all the included features is given in Appendices A and B.

|  | Data set 1 (DS1) | Data set 2 (DS2) |
| --- | --- | --- |
| | Average ± SD [range] | Average ± SD [range] |
| **No. of cows included** | **3754** | **2075** |
| **Theoretical 305-day yield** | 8261 ± 1772 [2688 ; 13755] | 8317 ± 1874 [2689 ; 13755] |
| **RMSE<sup>1</sup> [kg.day<sup>-1</sup>]** | 3.4 ± 1.4 [1 ; 28] | 3.8 ± 1.5 [1.2 ; 28] |
| **Peak yield [kg]** | 34 ± 6 [15 ; 61] | 34 ± 6 [14 ; 61] |
| **Days in milk of peak lactation [day]** | 59 ± 25 [1 ; 248] | 63 ± 27 [1 ; 207] |
| **Rate increasing part of the lactation curve [kg.day<sup>-1</sup>]** | 0.64 ± 0.28 [0.08 ; 6.7] | 0.62 ± 0.31 [0.08 ; 6.7] |
| **Persistency of the lactation curve [kg.day<sup>-1</sup>]** | -0.05 ± 0.03 [-0.15 ; 0.12] | -0.04 ± 0.03 [-0.15 ; 0.12] |
| **No. of perturbations per lactation** | 6.0 ± 3.1 [0 ; 17] | 6.3 ± 3.2 [0 ; 17] |
| **Losses in perturbations per lactation [kg]** | 148 ± 124 [0 ; 1137] | 154 ± 132 [0 ; 1129] |
| **Activity skewness** | | -0.05 ± 1.64 [-5.19 ; 8.77] |
| **No. of sharp and short activity peaks** | | 3.9 ± 2.7 [0 ; 18] |
| **Days in milk of first activity peak [days]** | | 56 ± 64 [1 ; 305] |
| **No. of general activity changes** | | 24 ± 16 [2 ; 94] |

<sup>1</sup> RMSE: Root mean squared error of the Wood model fitted on each lactation curve

Both the milk yield and the activity features were defined using biological knowledge of how these time series typically respond to changes in health and nutritional status (Højsgaard and Friggens, 2010; Codrea et al., 2011; Bjerre-Harpoth et al., 2012). We thereby always started from the data-own baseline, which was identified using a median smoother that is insensitive to sudden changes or differences in absolute value or baselines. Through characterization of the residuals from this median smoother and the correction and standardization at herd level, group changes and differences in the intensity of the responses between herds were corrected for.

### Predicting Lifetime Resilience Ranking from Milk Yield Features

#### Pearson Linear Correlations

The Pearson linear correlation coefficients between the lactation SF and RR are shown in Figure 3. A high (positive or negative) correlation between a SF and the RR suggests that there is a large effect of the SF on the RR. The average correlations per SF over all farms varied between *ρ* = −0.134 and *ρ* = +0.1492, and for some of the farms individual features showed correlations of more than +/- 0.4. Visual exploration of the correlation scatter plots (results not shown) did not show non-linear relationships either. Although consistency would have been expected, Figure 3 already suggests that there is only little consistency across farms in which SF can be predictive for the lifetime RR of the cows of a particular farm. For example, some of the features represent the effect of health events on the milk yield data through the characterization of perturbations. Because one could imagine that severe health events and the associated losses negatively affect longevity on all the farms, we supposed that there would be a consistently high correlation between perturbation-related SF and the RR. However, this consistency is lacking, especially in terms of the correlations’ sign (negative vs. positive). The highest and most consistent correlations are obtained for SF representing model fit and size of the residuals (RMSE of the Wood model (SF2), number of residuals below 85% of the predicted value (SF23) and average size of the 3 largest negative residuals (SF24)).

**Figure 3.**
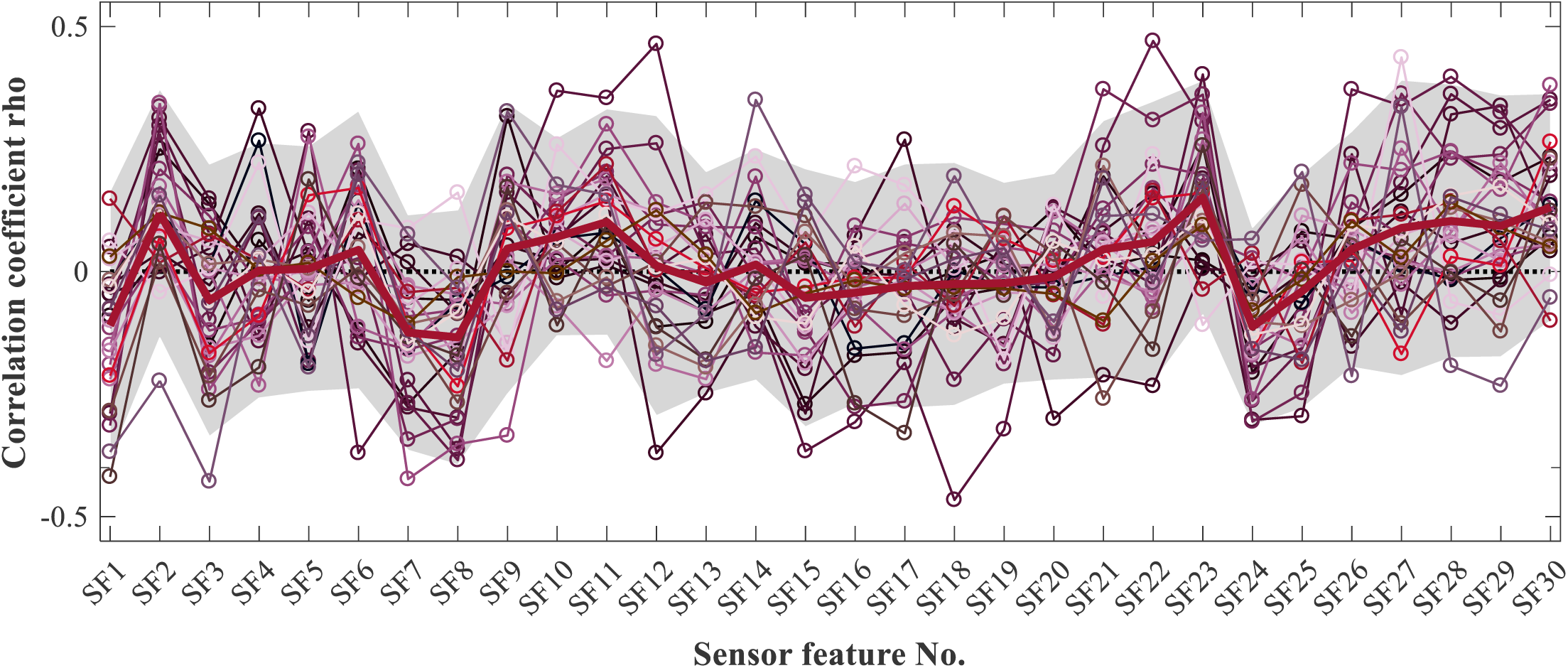
Pearson linear correlation coefficients between the lifetime resilience rank at farm level and the 30 lactation sensor features (**SF**) calculated on first parity data. Each individual thin line represents the correlations for a particular farm (total No. = 27). The shading represents the 95% confidence interval on the correlation coefficients over all farms. The details of the SF are described in Appendix A. The lack of consistency in the correlations’ signs and the magnitude demonstrate the large variability in the relation between the lifetime resilience and the lactation SF.

#### Model calibration

In the stepwise procedure for selecting the SF that are included in the multilinear regression model, the best possible combination of SF to fit the RR on each farm is determined, and SF are included or excluded dependent on whether they improve the model fit. The cross-validation step reveals whether the final model structure overfitted the training data and whether the selected SF are indeed meaningful for predicting the RR. If a SF is included in the multilinear regression model, the absolute value of its regression coefficient is directly related to its effect because of the standardization procedure of the SF (mean centering and equalizing the variance).

The *R*^*2*^_*adj*_ of the individual multivariate linear regression models of each farm varied between 0.03 and 0.61 (0.22 ± 0.16, mean ± SD) and the RMSE_TR_ was between 0.17 and 0.27 (0.23 ± 0.03, mean ± SD). Between 2 to 12 milk yield features were retained, and all SF were included at least once in one of the models of the individual farms (Table 3). The SF most often retained in the models were associated with the goodness-of-fit of the estimated Wood curves (SF21, 22, 25, 27, 28, 29), the size and number of perturbations (SF13, 14, 15), and their associated milk losses (SF11, 12, 18). This suggests that the SF that are proxies for (subclinical or chronic) health events are most informative over the RR, and thus these health events influence the cow’s longevity. It appears that farmers do take the health of the cows into account when making culling and re-insemination decisions, although not consistently and not on all farms. Moreover, the effects found in this study are rather weak, and in some cases it can be assumed that only the combined result of different features or only the extreme values might influence productive lifespan and RR. For example, a cow with severe clinical mastitis is likely to show a large and sudden drop in milk yield (Rajala-Schultz et al., 1999; Gröhn et al., 2004; Andersen et al., 2011) while a cow with subclinical mastitis may have a lactation curve which can be well modelled with a lactation model and which appears to be normal. Both can have impact on reproduction performance and the probability of culling (Lavon et al., 2010; Wathes, 2012; Wolfenson et al., 2015), but mathematically capturing the differences without additional health or treatment information is not possible. General lactation curve characteristics such as peak height and peak DIM, slopes, rate of the increasing phase of the lactation and persistency of the lactation after the peak) were only included in the models of only 12 out of 27 farms, suggesting that for example having a high milk production in the first lactation compared to herd mates barely affects the RR (and thus the ability to recalve and longevity). The variability between farms is demonstrated again by the fact that the regression coefficients were consistently above or below zero for only 2 out of the 30 SF, and so only 2 SF had a consistently positive or negative effect on the RR.

**Table 3.** Prediction results of the individual stepwise regression models in calibration and 10-times cross-validation (**CV**)

| Calibration, milk yield features only |  |  |  |  |  | Calibration, milk yield & activity features |  |  |  |  |  |  |  | CV<sup>8</sup>, milk yield features only |  |  |  | CV, milk yield & activity features |  |  |  |
| --- | --- | --- | --- | --- | --- | --- | --- | --- | --- | --- | --- | --- | --- | --- | --- | --- | --- | --- | --- | --- | --- |
| FARM ID | R<sup>2</sup><sub>adj</sub> | N<sup>1</sup> | N<sub>LAC</sub><sup>2</sup> | N<sub>GOF</sub><sup>3</sup> | N<sub>PERT</sub><sup>4</sup> | R<sup>2</sup><sub>adj</sub> | N | N<sub>LAC</sub> | N<sub>GOF</sub> | N<sub>PERT</sub> | N<sub>GEN</sub><sup>5</sup> | N<sub>L1</sub><sup>6</sup> | N<sub>L2</sub><sup>7</sup> | RMSE<sub>TR</sub><sup>9</sup> | RMSE<sub>CV</sub><sup>10</sup> | P<sub>C</sub><sup>11</sup> | P<sub>O</sub><sup>12</sup> | RMSE<sub>TR</sub> | RMSE<sub>CV</sub> | P<sub>C</sub> | P<sub>O</sub> |
| FARM1 | 0.10 | 8 | 2 | 4 | 2 |  |  |  |  |  |  |  |  | 0.26 | 0.27 | 45.9 | 3.7 |  |  |  |  |
| FARM2 | 0.16 | 5 | 0 | 2 | 3 |  |  |  |  |  |  |  |  | 0.24 | 0.27 | 44.3 | 1.1 |  |  |  |  |
| FARM3 | 0.05 | 4 | 1 | 1 | 2 |  |  |  |  |  |  |  |  | 0.25 | 0.27 | 41.5 | 3.6 |  |  |  |  |
| FARM4 | 0.27 | 3 | 0 | 1 | 2 |  |  |  |  |  |  |  |  | 0.21 | 0.21 | 50.0 | 0.0 |  |  |  |  |
| FARM5 | 0.29 | 8 | 0 | 2 | 6 |  |  |  |  |  |  |  |  | 0.22 | 0.25 | 45.8 | 6.8 |  |  |  |  |
| FARM6 | 0.53 | 5 | 1 | 1 | 3 |  |  |  |  |  |  |  |  | 0.17 | 0.18 | 70.0 | 0.0 |  |  |  |  |
| FARM7 | 0.31 | 4 | 0 | 2 | 2 |  |  |  |  |  |  |  |  | 0.23 | 0.24 | 59.0 | 2.6 |  |  |  |  |
| FARM8 | 0.42 | 11 | 2 | 5 | 4 |  |  |  |  |  |  |  |  | 0.19 | 0.23 | 45.6 | 6.3 |  |  |  |  |
| FARM9 | 0.08 | 2 | 0 | 1 | 1 |  |  |  |  |  |  |  |  | 0.25 | 0.25 | 44.6 | 2.2 |  |  |  |  |
| FARM10 | 0.16 | 8 | 2 | 2 | 4 |  |  |  |  |  |  |  |  | 0.24 | 0.27 | 48.9 | 5.6 |  |  |  |  |
| FARM11 | 0.14 | 7 | 2 | 0 | 5 |  |  |  |  |  |  |  |  | 0.24 | 0.26 | 37.1 | 3.4 |  |  |  |  |
| FARM12 | 0.40 | 9 | 1 | 3 | 5 | 0.51 | 8 | 1 | 2 | 2 | 1 | 2 | 0 | 0.20 | 0.24 | 55.9 | 1.7 | 0.18 | 0.20 | 52.5 | 1.7 |
| FARM13 | 0.21 | 8 | 2 | 3 | 3 | 0.32 | 7 | 1 | 3 | 0 | 1 | 2 | 0 | 0.23 | 0.25 | 45.0 | 11.2 | 0.21 | 0.22 | 52.6 | 1.0 |
| FARM14 | 0.28 | 6 | 1 | 3 | 2 |  |  |  |  |  |  |  |  | 0.21 | 0.23 | 45.6 | 1.3 |  |  |  |  |
| FARM15 | 0.06 | 3 | 2 | 0 | 1 |  |  |  |  |  |  |  |  | 0.19 | 0.19 | 41.8 | 1.8 |  |  |  |  |
| FARM16 | 0.10 | 9 | 3 | 2 | 4 | 0.30 | 15 | 3 | 2 | 2 | 1 | 4 | 3 | 0.23 | 0.24 | 44.8 | 7.4 | 0.20 | 0.21 | 51.5 | 1.8 |
| FARM17 | 0.08 | 6 | 1 | 2 | 3 | 0.27 | 14 | 1 | 3 | 5 | 2 | 2 | 1 | 0.26 | 0.27 | 35.8 | 6.1 | 0.23 | 0.26 | 45.3 | 4.0 |
| FARM18 | 0.53 | 7 | 2 | 2 | 3 | 0.76 | 13 | 0 | 3 | 6 | 2 | 1 | 1 | 0.16 | 0.19 | 55.0 | 5.0 | 0.10 | 0.15 | 77.5 | 0.0 |
| FARM19 | 0.18 | 6 | 3 | 1 | 2 | 0.34 | 11 | 2 | 4 | 2 | 0 | 2 | 1 | 0.23 | 0.25 | 55.0 | 8.3 | 0.19 | 0.24 | 52.3 | 1.5 |
| FARM20 | 0.14 | 5 | 0 | 2 | 3 |  |  |  |  |  |  |  |  | 0.24 | 0.25 | 40.7 | 5.0 |  |  |  |  |
| FARM21 | 0.21 | 4 | 1 | 1 | 2 | 0.33 | 5 | 0 | 1 | 1 | 1 | 0 | 2 | 0.22 | 0.22 | 35.9 | 5.3 | 0.20 | 0.22 | 43.5 | 0.0 |
| FARM22 | 0.30 | 6 | 1 | 0 | 5 | 0.67 | 23 | 3 | 5 | 9 | 2 | 2 | 2 | 0.18 | 0.20 | 45.6 | 1.4 | 0.11 | 0.17 | 66.2 | 0.0 |
| FARM23 | 0.11 | 5 | 0 | 1 | 4 | 0.24 | 14 | 0 | 1 | 4 | 1 | 4 | 4 | 0.26 | 0.26 | 39.3 | 2.8 | 0.23 | 0.25 | 44.4 | 6.7 |
| FARM24 | 0.08 | 5 | 1 | 1 | 3 | 0.36 | 13 | 1 | 3 | 1 | 2 | 4 | 2 | 0.26 | 0.26 | 43.4 | 5.9 | 0.21 | 0.22 | 53.2 | 0.5 |
| FARM25 | 0.03 | 3 | 0 | 2 | 1 | 0.20 | 12 | 1 | 4 | 1 | 2 | 2 | 2 | 0.26 | 0.26 | 41.9 | 16.5 | 0.23 | 0.24 | 49.4 | 5.2 |
| FARM26 | 0.61 | 12 | 1 | 2 | 9 | 0.62 | 8 | 1 | 1 | 4 | 1 | 1 | 0 | 0.15 | 0.21 | 62.0 | 2.0 | 0.16 | 0.18 | 84.0 | 4.2 |
| FARM27 | 0.15 | 6 | 1 | 3 | 2 | 0.32 | 13 | 2 | 2 | 5 | 0 | 3 | 1 | 0.26 | 0.26 | 40.4 | 2.5 | 0.22 | 0.24 | 48.5 | 3.7 |

<sup>1</sup> N: Total number of sensor features retained in the stepwise linear regression;
<sup>2</sup> N<sub>LAC</sub>: Number of features linked with the milk yield shape characteristics retained in the final regression models;
<sup>3</sup> N<sub>GOF</sub>: Number of features linked with the goodness-of-fit of the iterated Wood curve fitted on the lactation data retained in the final models;
<sup>4</sup> N<sub>PERT</sub>: Number of features linked with the identified perturbations retained in the final models;
<sup>5</sup> N<sub>GEN</sub>: Number of features linked with the general activity time series retained in the final models;
<sup>6</sup> N<sub>L1</sub>: Number of features linked with the activity peaks assumed to relate to estrus behavior retained in the final models;
<sup>7</sup> N<sub>L2</sub>: Number of features linked with periods of high or low average activity assumed to relate to general deviation in behavior or activity level in the final models;
<sup>8</sup> CV: Results of the 10-fold cross-validation
<sup>9</sup> RMSE<sub>TR</sub>: Average root mean squared error of the fitted scaled lifetime resilience ranking of the cows in the training set;
<sup>10</sup> RMSE<sub>CV</sub>: Average root mean squared error of cross-validation for the predicted scaled lifetime resilience ranking of the cows in the validation set;
<sup>11</sup> P<sub>C</sub>: Proportion of the animals on each farm predicted in the correct category (low, **L**, moderate, **M** or high, **H**);
<sup>12</sup> P<sub>O</sub>: Proportion of the animals on each farm predicted in the opposite category: cows predicted as being in the lowest third while being in the highest third and vice versa

#### Cross validation

With the CV, the repeatability, generality and overfitting of the models was tested. The within-farm CV showed similar performances (RMSEC = 0.24 ± 0.03) compared to the RMSE_TR_ of the initial models with all animals included, indicating that models were not overfitted. On average 46.7 ± 8.0% of the animals were classified in the correct H, M, L category, and on average 4.4 ± 3.5% of the cows were classified ‘high’ where they should have been ‘low’ and vice versa. Also here the large differences in model performance between the different farms stands out, with a range of correctly classified cows between35.8 and 70.0%, and the range of oppositely classified cows between 0.0 and 16.5%. When looking deeper into which cows are predicted in the right category, it was found that the models do not solely predict animals exiting the herd after the first lactation correctly but also cows exiting the herd in a later lactation. For cows culled already in the first lactation, health issues are expected to be the major reason (Pinedo et al., 2010). For cows culled at a later stage, it might not be expected that first lactation features are predictive for the RR unless these characteristics are highly repeatable over time or represent chronic or repeating conditions that fail to cure. However, information on the exact reason and timing of culling of the cows could not be taken into account in the model as this information was not available in the data sets. As it is expected that this information can contribute to the model, future research should focus on solving this multidimensionality issue and developing new ways to take these complex interactions into account.

### Predicting Lifetime Resilience Ranking from both Yield and Activity Features

#### Pearson Linear Correlations

Data set 2 consisted of 13 farms for which, besides milk yield features (No. = 30 features), activity features (No. = 15 features) could also be calculated. These features included both general characteristics of the daily activity (skewness, variability, absolute daily level) and specific features associated with short term and longer period activity changes. Farm-individual Pearson correlation coefficients (Figure 4) between the activity features and the RR varied between *ρ* = −0.41 and *ρ* = +0.44 and only the number of short activity peaks in (SF34) was consistently associated with a higher ranking (lower number of peaks is associated with a higher resilience, *ρ* = 0.29 ± 0.10, [0.16; 0.44]). Several other activity features also had correlations nearly consistently above or below zero, but these correlations stayed relatively low on average.

**Figure 4.**
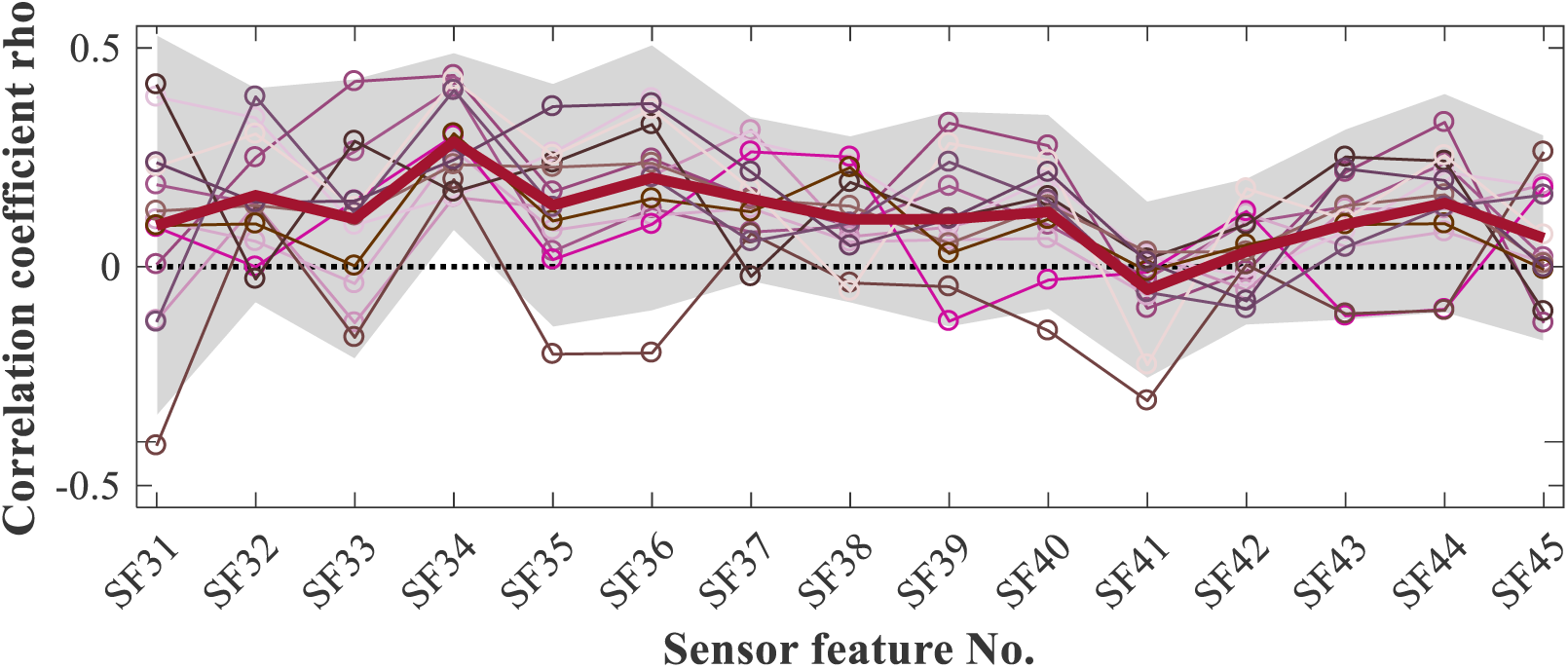
Pearson linear correlation coefficients between the lifetime resilience rank at farm level and the 15 activity sensor features (**SF**) calculated on first parity data. Each individual thin line represents the correlations of a particular farm (total No. = 13). The shading represents the 95% confidence interval on the correlation coefficients over all farms. The details of the activity SF are described in Appendix B. The lack of consistency in the correlations’ signs and the magnitude demonstrate the large variability in the relation between the lifetime resilience and the activity SF. Only SF34 (i.e., number of sharp activity peaks corresponding to estrus) have a consistently positive correlation with the lifetime resilience rank.

#### Model Calibration

The stepwise linear regression models included 6 to 24 SF (both activity and milk yield features) and had *R*^*2*^_*adj*_ values between 0.2 and 0.76 and RMSE_TR_ values between 0.128 and 0.24. The number of activity features retained in the final models was between 2 and 10, so the activity sensors seemed to be of added value for all of the farms in predicting their RR. Including activity features gave a higher *R*^*2*^_*adj*_ and a lower RMSE_TR_ in the calibration, while the number of features retained was sometimes higher and sometimes lower. Again, there was very little consistency over the different farms in which features were included in the final models. None of the SF was kept in the model of all farms. The number of activity peaks and DIM of the first peak were retained most often (respectively 8 and 11 out of 13 times) and with a consistently positive regression coefficient (respectively 0.036 to 0.122 and 0.042 to 0.134). Three of the SF (6.6%) were never retained in any of the individual farm models.

#### Cross validation

The CV, using the same cross-validation sets for these farms as in DS1 (i.e., the same animals were included in each set), showed reasonable performance, with an RMSECV of 0.22 ± 0.03 (range 0.15 to 0.26). On average 55.5 ± 12.1% (range 43.5 to 84.0%) of the cows were predicted in the correct category (H, M or L) and 2.3 ± 2.1% (range 0.0 to 6.7%) of them were predicted ‘high’ where they were actually ‘low’ or vice versa. This means that from the wrongly classified animals respectively 91.8% and 94.1% were predicted in an adjacent category. Over all the farms, including activity features improved the correct classification with 9.3 ± 7.9% (*P* < 0.01). The classification worsened in only two farms compared to when only milk yield features were included. The proportion classified in the opposite category decreased with on average 3.5 ± 4.5%, ranging from 5.9% to 2.3% (significant difference, *P* < 0.01).

Despite the variability between and within farms and the fact that we could not find SF that were commonly informative to predict RR over all farms, the prediction and classification performance of the individual farm models was in many cases significantly higher than the product of a random classification (i.e., a third correctly classified). Furthermore, including the activity features demonstrated a significant added value compared with using the daily milk yield features alone (*P* < 0.01). A correct classification of up to 84% of the animals suggests that at least part of the variability in the RR is correctly captured by the SF.

One way to explain the lack of a common model structure and the observed differences in prediction performance is the variability in culling, reproduction and health management between farms and even within farms. For example, management practices might have differed over the considered time span because of changing motivations and preferences of the farm staff, the economic context, the animals’ genotypes and phenotypes, the farm facilities, the feed etc. Besides the time-varying component, the following factors can also explain part of the limited prediction performance on some farms: (1) for this study, only features of the first lactation were included to ensure applicability of the model for decision support; (2) the lifetime resilience ranking is based on the limited data available in the commercial situation and was defined by experts; (3) there is a large difference in the number of animals included per farm, possibly affecting the results.

### Model Use and Implications

#### Predicting Lifetime Resilience Ranking Supports Breeding and Culling Decisions

Reliable prediction of lifetime resilience within a farm would allow for a more consistent approach to the management actions concerning advanced breeding (e.g., sexed semen, embryo transfer, ovum pick-up, use of beef semen, or selection of animals not to breed the replacement heifers from) or culling decisions after the first lactation (Mapletoft and Hasler, 2002; Vandeweerd et al., 2012; Boichard et al., 2015). In this way, breeding decisions for cows in the second parity and higher could be taken using both the genetic or genomic (available once the animal is born) and the phenotypic sensor-derived information (once an animal completed her first lactation). The latter would provide information on how well the animal performs in her specific farm environment, which optimizes sustainable productivity from the *available* animals on farm. In practice, discrimination between cows with high and low lifetime resilience would benefit a farmer even when the exact rankings remain unknown because farmer’s decision would not generally be different for e.g., the 5^th^ or the 10^th^ ranked cow in the herd. Moreover, prediction of lifetime resilience also allows for the identification of animals with a low expected RR. These cows can be targeted for more detailed monitoring in higher lactations.

This study also advocates the need for evidence-based decision making on modern dairy farms supporting more economically-sound and sustainable management actions. Despite the high prediction performance on some of the farms, the lack of a common model structure and the low performance on other farms suggest that further data-based rationalization of decisions is needed. In order to do so, dedicated data processing in which the biology of the cows within their farm contexts is taken into account is essential, and ideally also other key farm context indicators should be included in the resulting tools (e.g., herd demographics, robot or parlor capacity, economic environment, etc.).

#### Model use beyond decision support

With the collection and re-evaluation of this sensor-based information over many years, also general phenotypic information on complex traits for future breeding goals is collected at herd level. From this, sires that perform well under many different environmental conditions can be identified. In this way, the proposed tool can be used in the context of precision phenotyping of traits that are the combined result of physiological well-being and performance, and future genetic selection based on these new traits becomes possible.

## CONCLUSIONS

In this study, we demonstrate that resilience ranking and productive lifespan of modern dairy cows on AMS farms in Belgium and the UK could be predicted using farm-individual models based on first lactation sensor data. With the milk yield and activity SF selected at farm level, we reached classification performances (low, moderate, high resilience) of up to 84%, while only 2.3 ± 2.1% (mean ± SD) of the cows were predicted in the opposite category. This shows the potential of high-frequency milk yield and activity sensor data to rationalize evidence-based breeding and culling decisions. However, a common model structure across all farms could not be found, which shows the variability between farms and which highlights the need for biologically sound and context-dependent data processing tools. Once a lifetime resilience-predicting tool is established, the farmer and the livestock sector could not only benefit from it for management and decision support, but also at genetic level in the context of new precision phenotyping proxies for complex traits.

## ACKNOWLEDGEMENTS

Ines Adriaens received funding from the Research Foundation Flanders through grants No. 11ZG916N and V423719N and from a KU Leuven postdoctoral mandate grant No. PDM/19/132. The data from this study were collected in the format of farm software back-up files by the authors and the resulting database used is owned by the authors. Part of the data were collected in the context of the VLAIO LA-trajectory ‘MastiMan’, grant No. HBC.2016.0774 by DVM Igor Van den Brulle and dr. DVM Sofie Piepers from the University of Ghent, Flanders (Department of Reproduction, Obstetrics and Herd Health, M-Team and Mastitis and Milk Quality Research Unit). This work is part of the GenTORE project that has received funding from the European Union’s Horizon 2020 research and innovation program under grant agreement No. 727213. Also Katherine Lumb (RAFT Solutions Ltd., Ripon, United Kingdom) contributed to the data collection. We thank dr. Carmen Adriaens (Bernstein Laboratory, MGH, Boston, United States) for her critical reading of the manuscript.

## APPENDICES

### Appendix A Definition and Calculation of Milk Yield Sensor Features

In Table A1, the details of the different sensor features (**SF**) calculated from daily milk yield data included in the prediction models are given. The Iterated Theoretical Wood model (**ITW**) is the result of the iterative fitting and refitting procedure excluding perturbations to estimate the shape of the theoretical lactation curve. All SF are standardized at herd level before entering them in the models, by 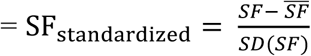 with SF each sensor feature, 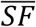 the average of each SF for a herd, and *SD(SF)* the standard deviation for that SF for a herd.

**Table A1.**
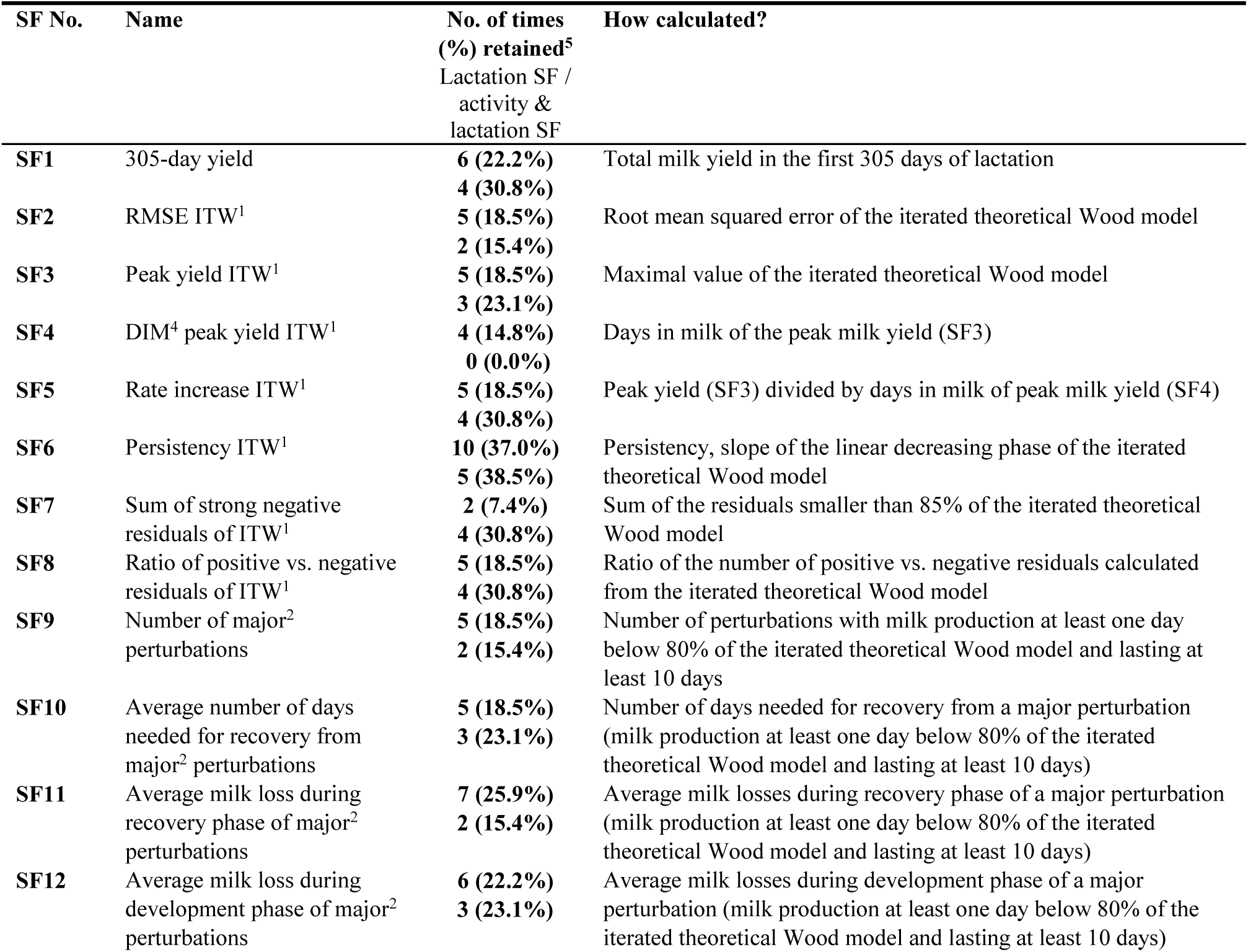
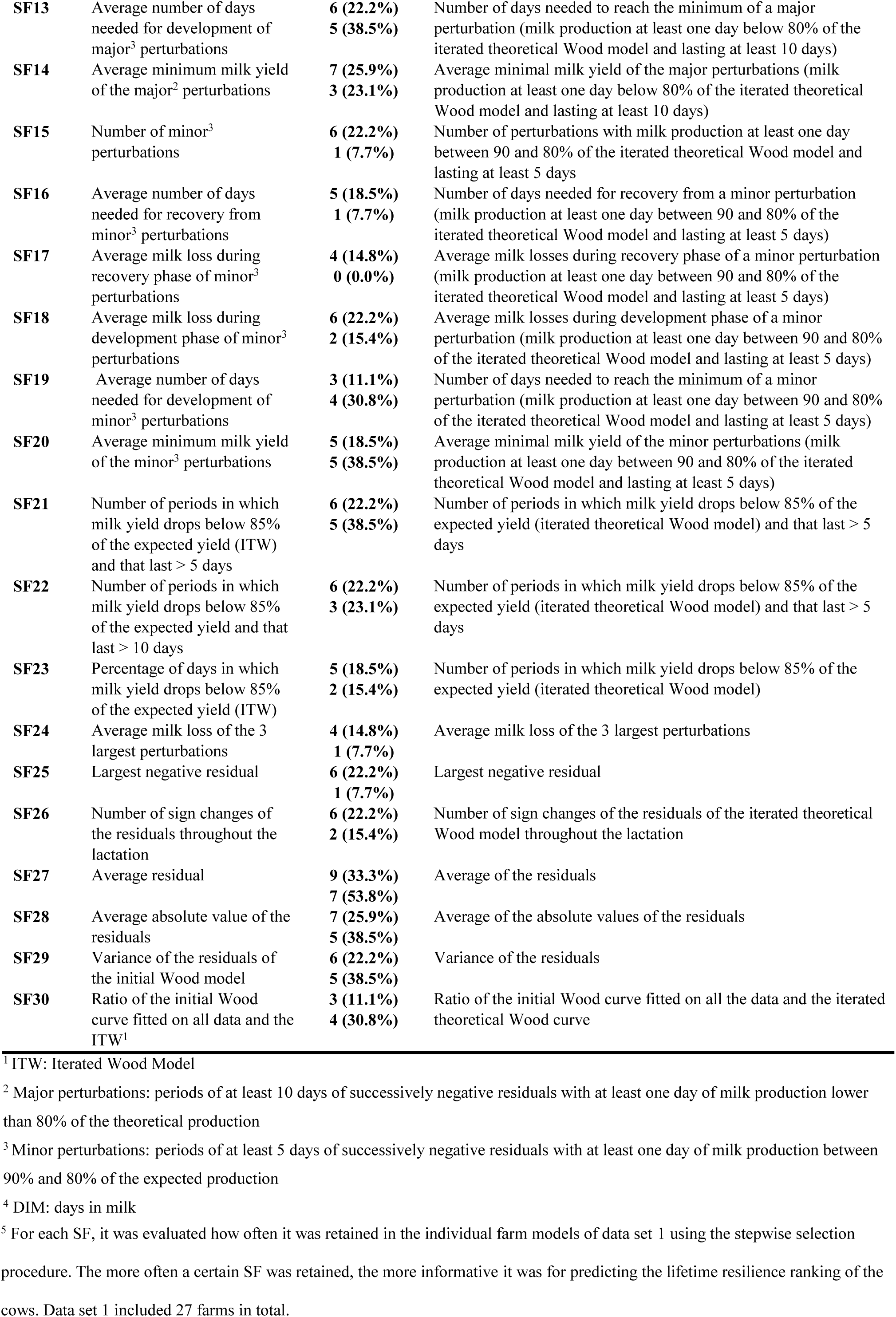
Lactation sensor features (**SF**) and their calculation included in the prediction models for lifetime resilience ranking

| SF No. | Name | No. of times<br>(%) retained<sup>5</sup><br>Lactation SF /<br>activity &<br>lactation SF | How calculated? |
| --- | --- | --- | --- |
| **SF1** | 305-day yield | **6 (22.2%)**<br>**4 (30.8%)** | Total milk yield in the first 305 days of lactation |
| **SF2** | RMSE ITW<sup>1</sup> | **5 (18.5%)**<br>**2 (15.4%)** | Root mean squared error of the iterated theoretical Wood model |
| **SF3** | Peak yield ITW<sup>1</sup> | **5 (18.5%)**<br>**3 (23.1%)** | Maximal value of the iterated theoretical Wood model |
| **SF4** | DIM<sup>4</sup> peak yield ITW<sup>1</sup> | **4 (14.8%)**<br>**0 (0.0%)** | Days in milk of the peak milk yield (SF3) |
| **SF5** | Rate increase ITW<sup>1</sup> | **5 (18.5%)**<br>**4 (30.8%)** | Peak yield (SF3) divided by days in milk of peak milk yield (SF4) |
| **SF6** | Persistency ITW<sup>1</sup> | **10 (37.0%)**<br>**5 (38.5%)** | Persistency, slope of the linear decreasing phase of the iterated theoretical Wood model |
| **SF7** | Sum of strong negative residuals of ITW<sup>1</sup> | **2 (7.4%)**<br>**4 (30.8%)** | Sum of the residuals smaller than 85% of the iterated theoretical Wood model |
| **SF8** | Ratio of positive vs. negative residuals of ITW<sup>1</sup> | **5 (18.5%)**<br>**4 (30.8%)** | Ratio of the number of positive vs. negative residuals calculated from the iterated theoretical Wood model |
| **SF9** | Number of major<sup>2</sup> perturbations | **5 (18.5%)**<br>**2 (15.4%)** | Number of perturbations with milk production at least one day below 80% of the iterated theoretical Wood model and lasting at least 10 days |
| **SF10** | Average number of days needed for recovery from major<sup>2</sup> perturbations | **5 (18.5%)**<br>**3 (23.1%)** | Number of days needed for recovery from a major perturbation (milk production at least one day below 80% of the iterated theoretical Wood model and lasting at least 10 days) |
| **SF11** | Average milk loss during recovery phase of major<sup>2</sup> perturbations | **7 (25.9%)**<br>**2 (15.4%)** | Average milk losses during recovery phase of a major perturbation (milk production at least one day below 80% of the iterated theoretical Wood model and lasting at least 10 days) |
| **SF12** | Average milk loss during development phase of major<sup>2</sup> perturbations | **6 (22.2%)**<br>**3 (23.1%)** | Average milk losses during development phase of a major perturbation (milk production at least one day below 80% of the iterated theoretical Wood model and lasting at least 10 days) |
| **SF13** | Average number of days needed for development of major<sup>3</sup> perturbations | **6 (22.2%)**<br>**5 (38.5%)** | Number of days needed to reach the minimum of a major perturbation (milk production at least one day below 80% of the iterated theoretical Wood model and lasting at least 10 days) |
| **SF14** | Average minimum milk yield of the major<sup>2</sup> perturbations | **7 (25.9%)**<br>**3 (23.1%)** | Average minimal milk yield of the major perturbations (milk production at least one day below 80% of the iterated theoretical Wood model and lasting at least 10 days) |
| **SF15** | Number of minor<sup>3</sup> perturbations | **6 (22.2%)**<br>**1 (7.7%)** | Number of perturbations with milk production at least one day between 90 and 80% of the iterated theoretical Wood model and lasting at least 5 days |
| **SF16** | Average number of days needed for recovery from minor<sup>3</sup> perturbations | **5 (18.5%)**<br>**1 (7.7%)** | Number of days needed for recovery from a minor perturbation (milk production at least one day between 90 and 80% of the iterated theoretical Wood model and lasting at least 5 days) |
| **SF17** | Average milk loss during recovery phase of minor<sup>3</sup> perturbations | **4 (14.8%)**<br>**0 (0.0%)** | Average milk losses during recovery phase of a minor perturbation (milk production at least one day between 90 and 80% of the iterated theoretical Wood model and lasting at least 5 days) |
| **SF18** | Average milk loss during development phase of minor<sup>3</sup> perturbations | **6 (22.2%)**<br>**2 (15.4%)** | Average milk losses during development phase of a minor perturbation (milk production at least one day between 90 and 80% of the iterated theoretical Wood model and lasting at least 5 days) |
| **SF19** | Average number of days needed for development of minor<sup>3</sup> perturbations | **3 (11.1%)**<br>**4 (30.8%)** | Number of days needed to reach the minimum of a minor perturbation (milk production at least one day between 90 and 80% of the iterated theoretical Wood model and lasting at least 5 days) |
| **SF20** | Average minimum milk yield of the minor<sup>3</sup> perturbations | **5 (18.5%)**<br>**5 (38.5%)** | Average minimal milk yield of the minor perturbations (milk production at least one day between 90 and 80% of the iterated theoretical Wood model and lasting at least 5 days) |
| **SF21** | Number of periods in which milk yield drops below 85% of the expected yield (ITW) and that last > 5 days | **6 (22.2%)**<br>**5 (38.5%)** | Number of periods in which milk yield drops below 85% of the expected yield (iterated theoretical Wood model) and that last > 5 days |
| **SF22** | Number of periods in which milk yield drops below 85% of the expected yield and that last > 10 days | **6 (22.2%)**<br>**3 (23.1%)** | Number of periods in which milk yield drops below 85% of the expected yield (iterated theoretical Wood model) and that last > 5 days |
| **SF23** | Percentage of days in which milk yield drops below 85% of the expected yield (ITW) | **5 (18.5%)**<br>**2 (15.4%)** | Number of periods in which milk yield drops below 85% of the expected yield (iterated theoretical Wood model) |
| **SF24** | Average milk loss of the 3 largest perturbations | **4 (14.8%)**<br>**1 (7.7%)** | Average milk loss of the 3 largest perturbations |
| **SF25** | Largest negative residual | **6 (22.2%)**<br>**1 (7.7%)** | Largest negative residual |
| **SF26** | Number of sign changes of the residuals throughout the lactation | **6 (22.2%)**<br>**2 (15.4%)** | Number of sign changes of the residuals of the iterated theoretical Wood model throughout the lactation |
| **SF27** | Average residual | **9 (33.3%)**<br>**7 (53.8%)** | Average of the residuals |
| **SF28** | Average absolute value of the residuals | **7 (25.9%)**<br>**5 (38.5%)** | Average of the absolute values of the residuals |
| **SF29** | Variance of the residuals of the initial Wood model | **6 (22.2%)**<br>**5 (38.5%)** | Variance of the residuals |
| **SF30** | Ratio of the initial Wood curve fitted on all data and the ITW<sup>1</sup> | **3 (11.1%)**<br>**4 (30.8%)** | Ratio of the initial Wood curve fitted on all the data and the iterated theoretical Wood curve |

<sup>1</sup> ITW: Iterated Wood Model
<sup>2</sup> Major perturbations: periods of at least 10 days of successively negative residuals with at least one day of milk production lower than 80% of the theoretical production
<sup>3</sup> Minor perturbations: periods of at least 5 days of successively negative residuals with at least one day of milk production between 90% and 80% of the expected production
<sup>4</sup> DIM: days in milk

### Appendix B. Definition and calculation of activity sensor features (SF)

First, the two-hourly activity data were aggregated in daily sums. Next, a moving median using a window of 4 days was calculated on these daily data time series to identify short spikes associated with estrous behavior (level 1; all spikes > 0.4* the maximal residual of the time series minus the moving median). A moving median of 20 days was calculated to identify periods with generally lower or higher activity (possibly associated with health events). A threshold of 20% of the minimal or maximal activity residuals was set to identify these altered activity period. The below explained calculated SF are based on the deviations from these median windows.

**Table B1.**
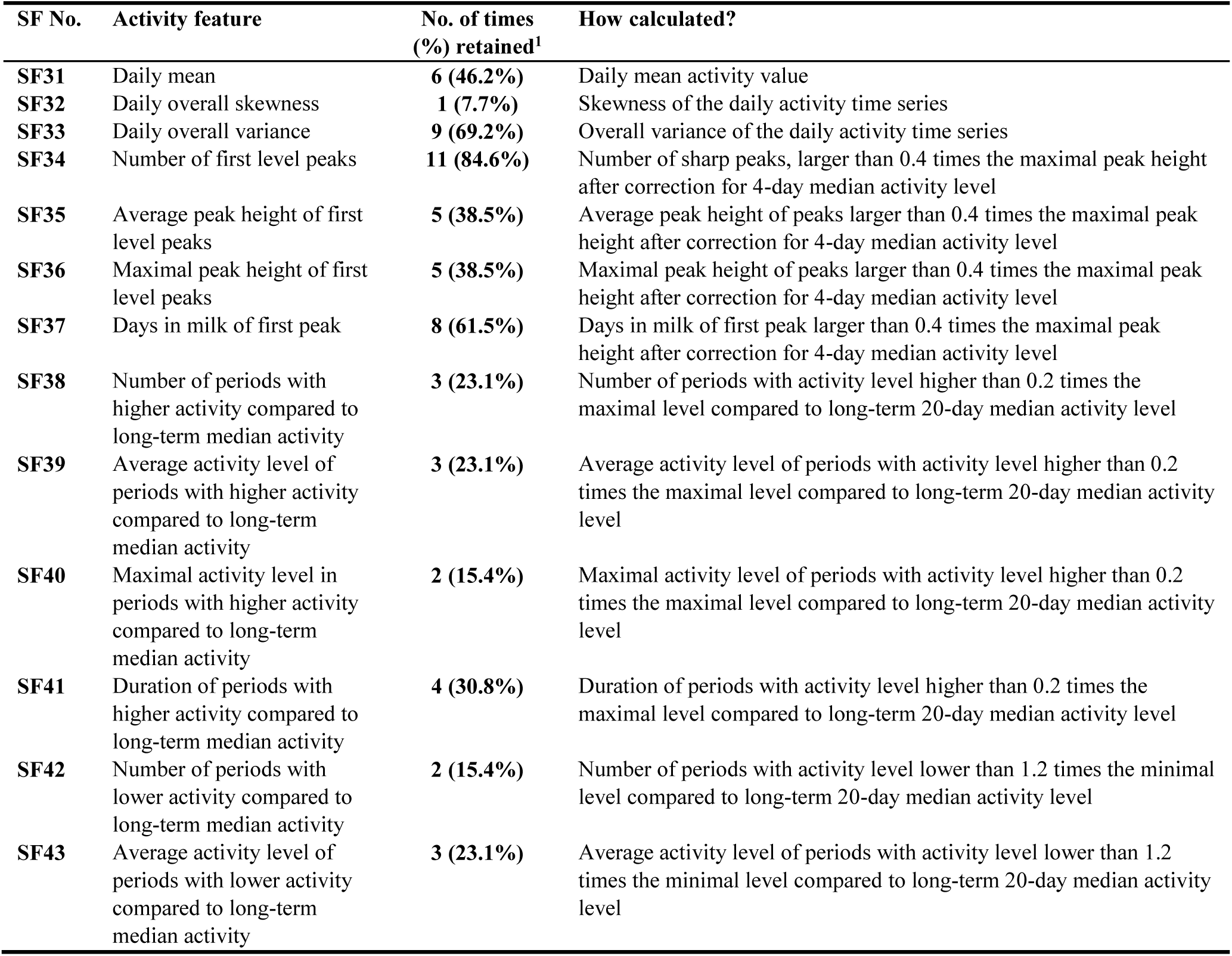
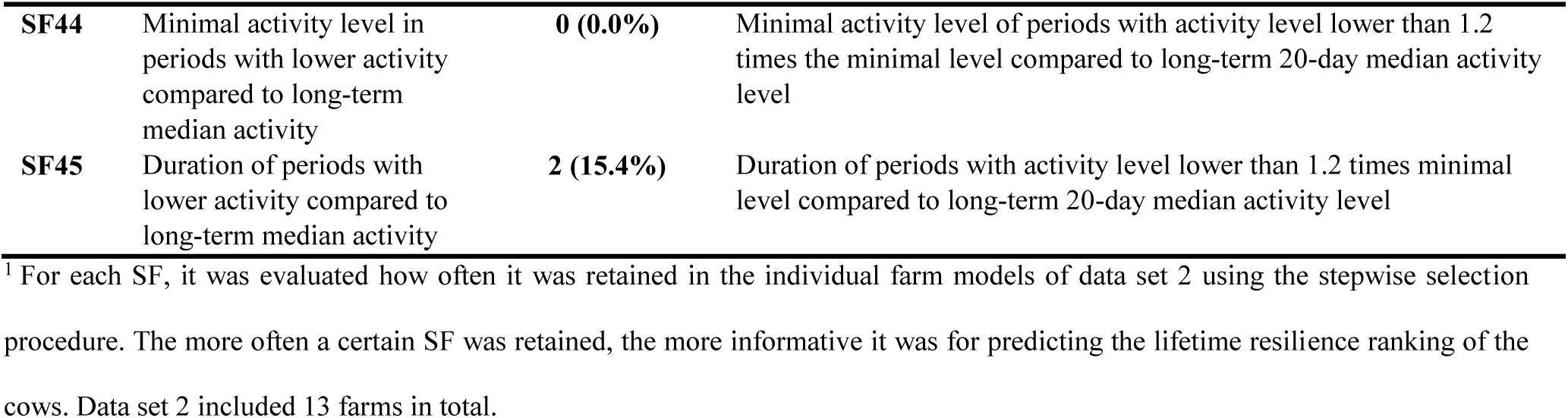
Activity sensor features (**SF**) and their calculation included in the prediction models for lifetime resilience ranking

| SF No. | Activity feature | No. of times (%) retained<sup>1</sup> | How calculated? |
| --- | --- | --- | --- |
| SF31 | Daily mean | 6 (46.2%) | Daily mean activity value |
| SF32 | Daily overall skewness | 1 (7.7%) | Skewness of the daily activity time series |
| SF33 | Daily overall variance | 9 (69.2%) | Overall variance of the daily activity time series |
| SF34 | Number of first level peaks | 11 (84.6%) | Number of sharp peaks, larger than 0.4 times the maximal peak height after correction for 4-day median activity level |
| SF35 | Average peak height of first level peaks | 5 (38.5%) | Average peak height of peaks larger than 0.4 times the maximal peak height after correction for 4-day median activity level |
| SF36 | Maximal peak height of first level peaks | 5 (38.5%) | Maximal peak height of peaks larger than 0.4 times the maximal peak height after correction for 4-day median activity level |
| SF37 | Days in milk of first peak | 8 (61.5%) | Days in milk of first peak larger than 0.4 times the maximal peak height after correction for 4-day median activity level |
| SF38 | Number of periods with higher activity compared to long-term median activity | 3 (23.1%) | Number of periods with activity level higher than 0.2 times the maximal level compared to long-term 20-day median activity level |
| SF39 | Average activity level of periods with higher activity compared to long-term median activity | 3 (23.1%) | Average activity level of periods with activity level higher than 0.2 times the maximal level compared to long-term 20-day median activity level |
| SF40 | Maximal activity level in periods with higher activity compared to long-term median activity | 2 (15.4%) | Maximal activity level of periods with activity level higher than 0.2 times the maximal level compared to long-term 20-day median activity level |
| SF41 | Duration of periods with higher activity compared to long-term median activity | 4 (30.8%) | Duration of periods with activity level higher than 0.2 times the maximal level compared to long-term 20-day median activity level |
| SF42 | Number of periods with lower activity compared to long-term median activity | 2 (15.4%) | Number of periods with activity level lower than 1.2 times the minimal level compared to long-term 20-day median activity level |
| SF43 | Average activity level of periods with lower activity compared to long-term median activity | 3 (23.1%) | Average activity level of periods with activity level lower than 1.2 times the minimal level compared to long-term 20-day median activity level |
| **SF44** | Minimal activity level in periods with lower activity compared to long-term median activity | **0 (0.0%)** | Minimal activity level of periods with activity level lower than 1.2 times the minimal level compared to long-term 20-day median activity level |
| **SF45** | Duration of periods with lower activity compared to long-term median activity | **2 (15.4%)** | Duration of periods with activity level lower than 1.2 times minimal level compared to long-term 20-day median activity level |

<sup>1</sup> For each SF, it was evaluated how often it was retained in the individual farm models of data set 2 using the stepwise selection procedure. The more often a certain SF was retained, the more informative it was for predicting the lifetime resilience ranking of the cows. Data set 2 included 13 farms in total.

## Notes

#### Summary of Updates

Additional explanations Revised introduction Revised and combined results and discussion section

